# Multiregional Representations of Intertemporal Decision Making in Human Single Neurons

**DOI:** 10.1101/2024.05.13.594032

**Authors:** Jay L. Gill, Mahmoud Omidbeigi, Jihye Ryu, Nanthia Suthana, Jonathan C. Kao, Ausaf Bari

**Affiliations:** Medical Scientist Training Program; University of California; Los Angeles, California; Department of Psychiatry and Biobehavioral Sciences, Jane and Terry Semel Institute for Neuroscience and Human Behavior; University of California; Los Angeles, California; Department of Neurosurgery; University of California; Los Angeles, California; Department of Psychology; University of California; Los Angeles, California; Department of Bioengineering; University of California; Los Angeles, California; Department of Electrical and Computer Engineering; University of California; Los Angeles, California

## Abstract

Characterization of the neural basis of delay discounting provides insight into the origin of impulsive decision-making, which underlies several psychiatric diseases including substance use disorder. Here, we identify human single unit representations of decisions, and their level of difficulty, in the orbitofrontal cortex, hippocampus and amygdala that are related to preferences for immediate vs. delayed rewards. Results provide an initial account of multiregional intracranial computations related to impulsive behaviors in humans.

## Main Text

Though patience is a virtue, humans generally prefer smaller, immediate rewards over larger, delayed rewards. A strong bias towards instant gratification results in heightened impulsivity— a core feature of addiction, suicide, and attention and impulse control related disorders^1^. Understanding the human intracranial neural activity that supports option appraisal, and how it is altered during impulsive decisions, is essential to the development of targeted therapeutics for individuals with pathologically risky behaviors.

Human neuroimaging and animal findings suggest that option contemplation and selection are highly distributed processes^2,3,4^. The orbitofrontal cortex (OFC) encodes subjective value and influences choice, the amygdala facilitates acquisition of stimulus-affect relationships^5,6^, and the hippocampus uses past experiences to qualify prospective outcomes that are distant in time^7^. The human single unit correlates of these functions remain largely uncharacterized, presenting a challenge in translating basic scientific findings to daily human decision behaviors that are often compromised in the setting of disease.

We present single unit data collected from the human OFC, amygdala, and hippocampus during performance of a delay discounting task where participants chose between a smaller, immediate and a larger, delayed reward. Using a logistic regression approach, we find that choice predictive neural activity is present throughout all three regions and is most strongly represented in the OFC. We also observe neural activity indicative of decision conflict between similarly valued options. Notably, units isolated in the OFC and hippocampus, but not the amygdala, were more predictive of choice compared to decision difficulty. Lastly, our findings demonstrate that hippocampal representations of choice and decision conflict are stronger in participants with slower temporal discounting of delayed rewards. These results are therefore an initial characterization of multiregional single unit activity related to intertemporal choice in humans.

Nine human participants with indwelling microelectrodes^8^ for epilepsy treatment performed a forced-choice delay discounting task (Figure 1 a-b; Supplementary Table 1; Methods). During each trial, participants were asked if they preferred $10 at a delay or an amount less than $10 immediately. Decisions across trials were used to quantify the unique rate at which each participant subjectively devalued delayed rewards, *k* (Methods; Figure 1c; Supplementary Figure 1). Larger discounting rates indicate a stronger preference for immediate over delayed rewards and have previously been related to increased impulsivity^1^. We then used the individual discounting rate to approximate the subjective value of the delayed offer and decision difficulty (i.e. the difference between the immediate offer value and the subjective value of the delayed reward) for each trial (Methods).

**Figure 1.**
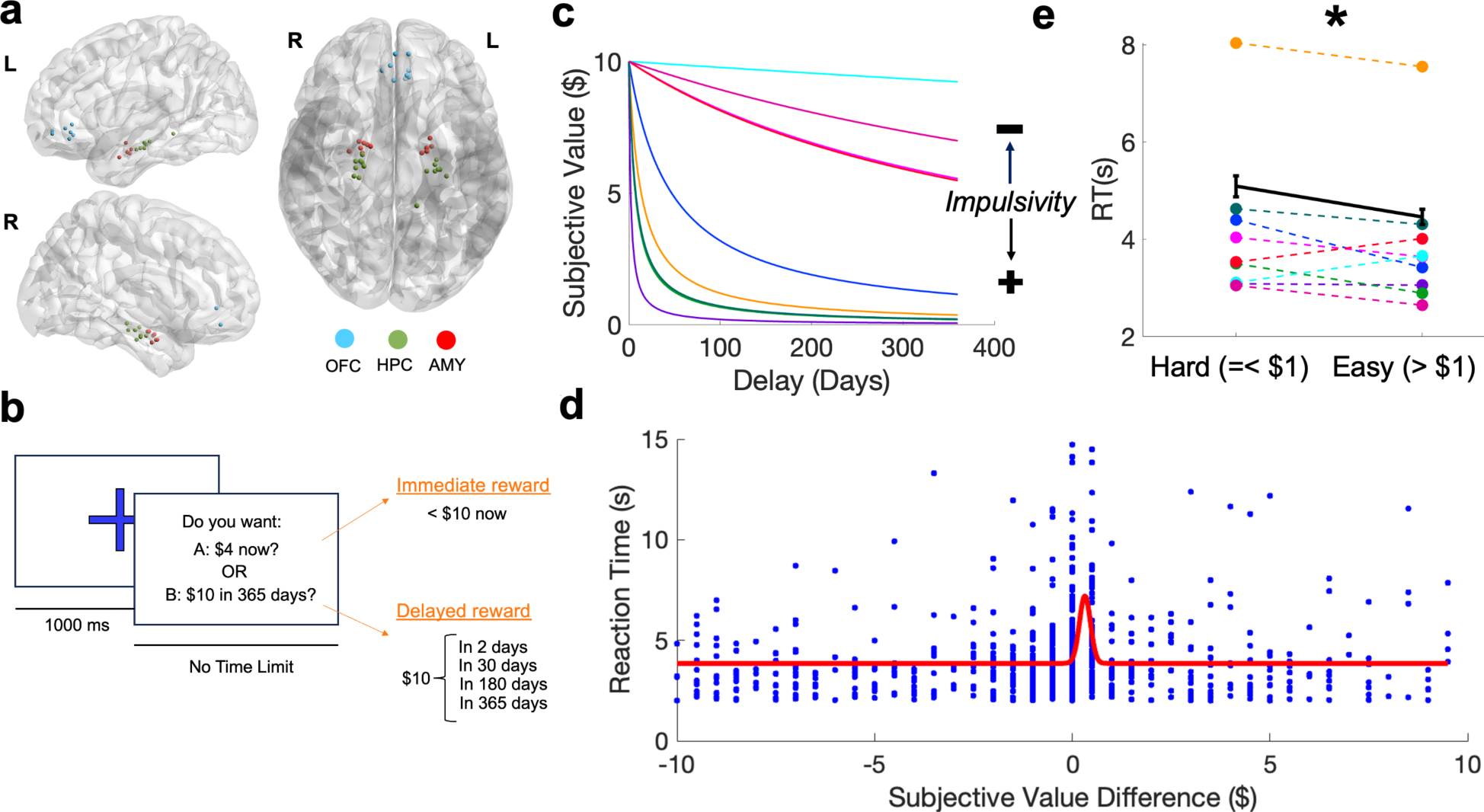
Individual Preferences and Proximity in Subjective Value of Offered Rewards Affect Intertemporal Choice. **a)** Sagittal (left [L]; right [R]) and axial views of microelectrodes from all participants localized in Montreal Neurological Institute (MNI) space. OFC: orbitofrontal cortex, AMY: amygdala, HPC: hippocampus. **b)** Schematic of delay discounting task. There was a 1000 millisecond (ms) fixation cross followed by a question prompt. During each trial, participants selected one of two options: (1) an immediate reward smaller than $10 (“A”) or (2) $10 at some fixed delay (“B”; “Delayed Reward”). **c)** Discounting curves for each participant (Supplementary Figure 1). A steeper slope indicates the individual more rapidly discounted delayed rewards and preferred smaller, immediate offers. **d)** Increased reaction time observed as participants decided between two options of similar subjective value. Individual points represent the difference in subjective value between offered options and the corresponding reaction time. The red line represents a Gaussian function fit to data using nonlinear regression^9^. **e)** Model parameters identified by the Gaussian function indicated that the rise in the curve (i.e. increase in reaction time) occurred surrounding a subjective value difference less than |$1|. |$1| was therefore used as the threshold for determining decision difficulty (hard vs. easy choice). Reaction time was significantly greater during hard compared to easy choices. * = p <0.05 using a linear effects model with a fixed effect of reaction time x subjective value and random effect of participant. Average reaction time for each subject during hard and easy choices are displayed using the same color code specified in **c.**

We first asked if decision difficulty was related to behavior across participants. Previous studies report longer reaction times (RTs) during decisions involving options with small (difficult) compared to large (easy) differences in subjective value (DSV). To quantify the relationship between DSV and RT, we fit a Gaussian function^9^ using nonlinear regression (Methods). This approach identified a rise in RT at $1.078 (Figure 1 e-d). Use of this amount as a threshold for defining easy (DSV >|$1|) vs. hard (DSV <|$1|) trials identified significantly longer RTs during difficult decisions (Figure 1e).

A total of 193 single units were isolated across participants (50 OFC, 68 Amygdala, 75 Hippocampus; Figure 1a). We first asked if neural activity differed when a participant was about to select an immediate vs. delayed option. To enhance our ability to compare activity across trials with uneven time lengths, analyses focused on activity during the peri-decision period (Methods). We used a logistic regression decoding approach (Methods; Figure 2c-d) and found 48 (96%), 51 (75%), and 51 (68%) units within the OFC, amygdala, and hippocampus, respectively, predicted choice prior to decision onset. We next asked if decision difficulty (hard [DSV =<|$1|] vs. easy [DSV >|$1|]) could similarly be predicted during the pre-decision period. We used the previously described decoding approach to predict whether a decision was easy or hard (Figure 2e). We identified 44 (88%), 57 (84%), and 64 (85%) units within the OFC, amygdala, and hippocampus, respectively, that exhibited significant decision difficulty encoding. Of the total number of choice and/or difficulty encoding neurons 84%, 71% and 60% of units in the OFC, amygdala, and hippocampus, respectively, exhibited encoding of both variables (Supplementary Figure 2). Though simultaneous peak encoding of both variables was infrequent, no significant temporal preferences (i.e., peak choice decoding preceding peak difficulty decoding) were observed across “both” encoding units (Supplementary Figure 3).

**Figure 2.**
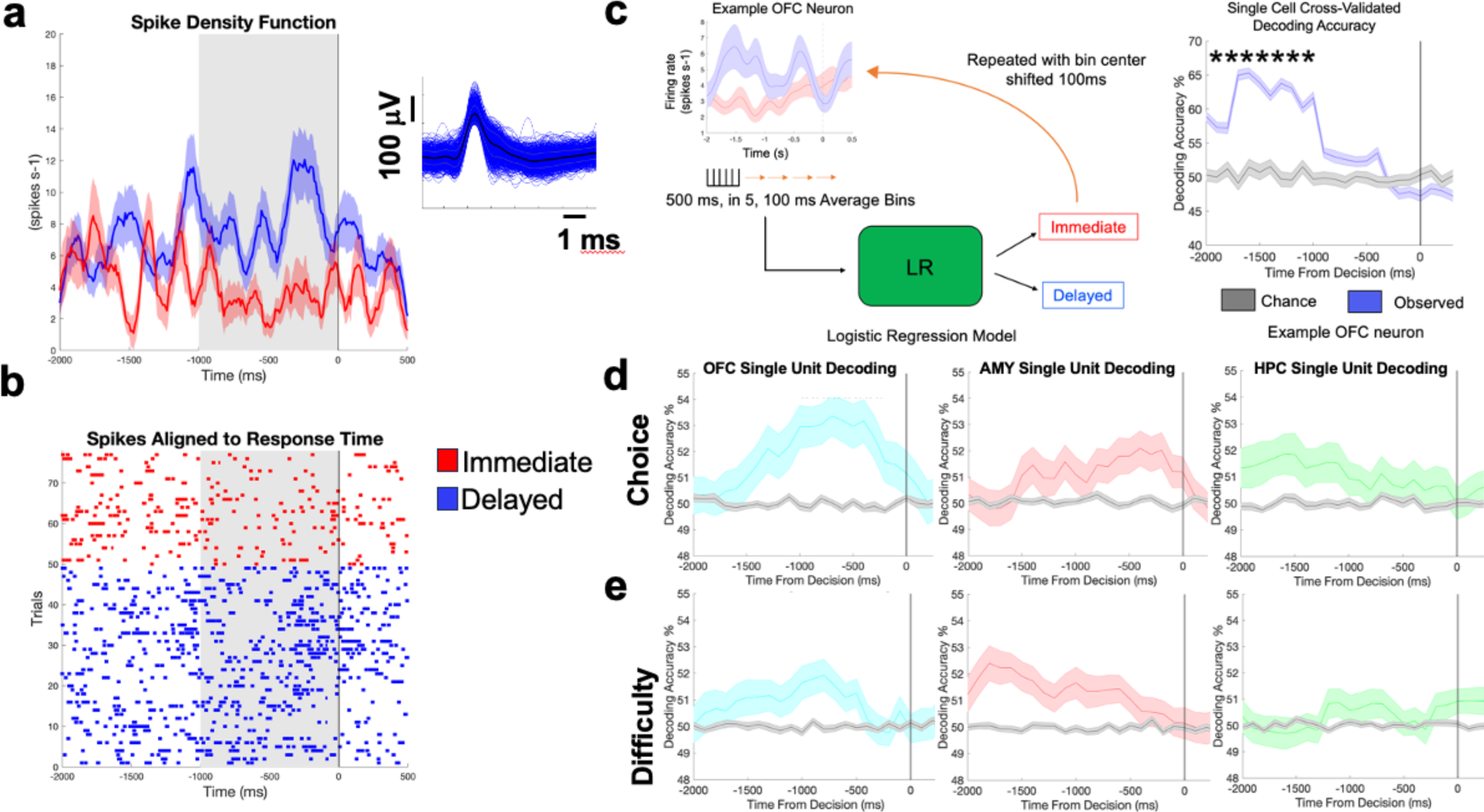
Intertemporal Choice is Reflected in Human Single Neurons in the Orbitofrontal Cortex, Amygdala, and Hippocampus. **a)** Spike density (average +/- standard error of mean [s.e.m.]) and raster plot **b)** for an example OFC unit with differential responses during immediate (red) and delayed (blue) choices. Average +/- standard error of all spike waveforms **a)** left. **c)** Schematic of single neuron decoding approach. During decoding, the firing rate of a neuron on a single trial was binned into 100 ms time bins. The firing rate was averaged across these 100ms. We then selected 5 bins (features) as the input to a logistic regression model. The model therefore saw 500ms of activity. At each time point the model predicted the choice made at the end of the corresponding trial (selection of an immediate or delayed reward). We then plotted decoding accuracy across time. The time on the x-axis corresponds to the absolute time of the midpoint of the 500ms input. A unit was called “choice encoding” if feeding the decoder activity at any point prior to the choice that led to an accuracy that was both above chance and greater than the decoding accuracy of a chance decoder trained on the same activity but shuffled labels (* = p<0.05; Bonferroni corrected). **d)** The average decoding accuracy +/- s.e.m of choice selective units from the OFC (left), amygdala (AMY; middle) and hippocampus (HPC; right). **e)** Same as **d** but for difficulty encoding units that were isolated using the same approach depicted in **c**, but with a decoder predicting whether a choice being made was hard or easy.

We next sought to characterize population-level encoding of choice and difficulty. A choice population decoder was built for each region using activity from units that exhibited significant choice encoding. Each model feature, N_1_, N_2_ … N_n_ (where n = number of choice selective units within a region across all participants; Methods), was the average firing rate of a cell in a 500 ms window centered around the time the unit exhibited its highest choice decoding accuracy. Models were trained on 50% of the available data using logistic regression and tested on the remaining 50% (Methods). This approach yielded population accuracies of 78%, 72.6%, and 73.8%, across the OFC, amygdala, and hippocampus respectively (Figure 3; Supplementary Figure 4-6). When this approach was used to decode difficulty, we found accuracies of 67% (OFC), 74% (amygdala), and 66% (hippocampus). Comparisons of average peak single cell choice vs. difficulty accuracies exhibited similar results where the OFC and hippocampus, but not the amygdala, exhibited significantly greater average peak single cell accuracies when decoding choice compared to difficulty (Figure 3).

**Figure 3.**
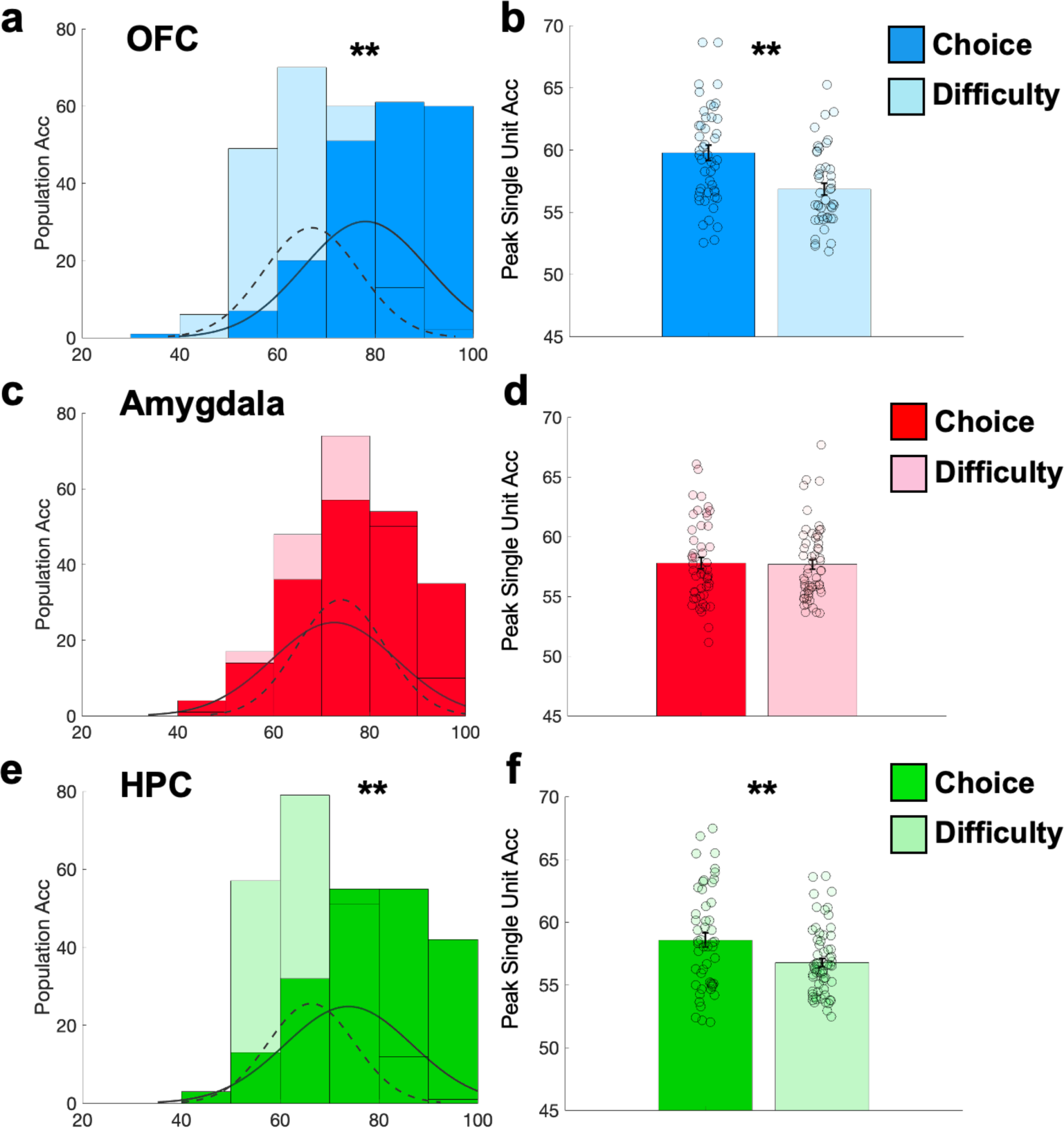
Peak Population and Single Neuron Differences in Choice and Decision Difficulty Encoding. (Right column) Histogram of decoding performance across cross validations from choice (darker shade of all colors, solid line) and difficulty (lighter shade of all colors, dashed line) decoders built using activity from units (features) that significantly encoded choice and difficulty, respectively, in the **a)** OFC (blue; choice mean +/- standard deviation [s.d.]: 78.00% +/- 12.72 s.d., difficulty mean +/- s.d.: 66.97% +/- 9.79; p < .001), **c)** amygdala (red; choice mean +/- standard deviation [s.d.]: 72.65% +/- 12.93 s.d., difficulty mean +/- s.d.: 74.07% +/- 9.09; p = .219), and **e)** HPC (green; choice mean +/- standard deviation [s.d.]: 73.85% +/- 12.86 s.d., difficulty mean +/- s.d.: 66.32 +/- 8.73; p < .001). * = p < 0.05, ** = p<0.01 using permutation testing. **(Left column)** Average peak single unit activity during decoding of choice (darker shade of all colors) and difficulty (lighter shade of all colors) in the **b)** OFC (blue; Choice: 48 units, mean = 59.76% +/- .61 s.e.m; Difficulty: 44 units, mean = 56.83% +/- .47 s.e.m; p <.001), **d)** amygdala (red; Choice: 51 units, mean = 57.79% +/- .48 s.e.m; Difficulty: 57 units, mean = 57.67% +/- .41 s.e.m; p = .687) and **e)** HPC (green; Choice: 51 units, mean = 58.56% +/- .56 s.e.m; Difficulty: 64 units, mean = 56.77% +/- .33 s.e.m; p = .004). * = p < .05, ** = p<.01 using linear mixed effect models.

To understand whether decoding dynamics were influenced by participant behavior, particularly their preference for immediate rewards, we constructed decoders using varying proportions of units from fast discounters (present favoring) and slow discounters (future favoring). Initially, we compared decoders using units exclusively from fast discounters with an equal number of units from all participants (to serve as a size-matched approximation of the initial behavior-agnostic population decoders; Methods). We observed differences between the “fast-only” and “all (fixed)” models in the hippocampus and amygdala, but not in the OFC (Supplementary Figure 7). Representation of choice was increased in “fast-only” compared to the “all (fixed)” model in the amygdala. Conversely, choice and difficulty representations were decreased in “fast-only” hippocampal models (Supplementary Figure 7). Further comparisons were made by building decoders exclusively from units of either fast or slow discounters, matched in number. We again observed reduced choice and difficulty representations in the hippocampus, while the amygdala showed increased choice representations in the “fast-only” models compared to the “slow-only” models (Supplementary Figure 8). Analysis of data at the single unit level did not reveal significant “slow” vs. “fast-only” hippocampal average peak accuracy differences (Supplementary Figure 9), however, amygdala “fast-only” isolated units exhibited a later decoding of choice and earlier decoding of difficulty compared to “slow-only” models (Supplementary Figure 10).

Decades of non-invasive human neuroimaging studies demonstrate that delay discounting behavior–related (subjective valuation of offered options and choice preferences) activity is reflected throughout a broad neural network that includes the OFC, amygdala, and hippocampus^3,4,10^. Though findings from animal models suggest that economic decisions are driven by single neuron representations of subjective value^11,12^, characterization of neural activity that supports the evaluation of distal, abstract (vs. appetitive) prospective outcomes in humans has been largely out of reach^13,14^. We begin to address this knowledge gap in the present study and demonstrate that 1) option appraisal (derived from approximations of participant specific subjective value of delayed rewards and behaviorally validated) and 2) choice are reflected in single neuron activity across the human amygdala, OFC, and hippocampus.

The importance of understanding the neural origins of delay discounting is underscored by its alteration across a myriad of disorders related to heightened impulsivity. Neuroimaging studies suggest that the rate at which individuals discount future rewards (i.e. the amount of preference given to the present vs. the future) is related to interactions between frontal and medial temporal regions driven by episodic thinking^2^. Notably, rats with hippocampal dysfunction demonstrate increased delay discounting^15,16^ and rats with amygdala lesions fail to learn cue-outcome associations^5^, highlighting the importance of MTL structures in acquisition of subjective value representations. Congruent findings are reported in humans where patients with amygdala damage fail to learn from aversive monetary outcomes^17^ and patients with MTL damage do not exhibit episodic thinking related reductions in temporal discounting^7^. Using models built with units from participants with fast vs. slow discounting behaviors, we demonstrate that the degree of single unit representation of task variables in the amygdala and hippocampus may underlie an individual’s preference for smaller, immediate vs. larger, delayed rewards.

We use a decoding approach to characterize neural activity discounting behaviors. This allowed us to isolate common neural dynamics across broad trial categories that were detected using traditional firing rate approaches. Though we comment on representations of delay discounting related variables/behaviors, how communication between discussed regions sculpts behavior remains uncharacterized and should be explored in future studies. Given the self-paced design, we focus analyses on peri-decision activity to characterize activity at identical periods across trials of varying lengths. This approach, which allowed us to effectively characterize the relationship between decision difficulty and reaction time, limits the ability to comment on state transitions leading up to decision making. Research in non-human primates demonstrates that value encoding neurons in OFC contributed to vacillating neural states during option contemplation that ultimately predicted choice^18^. Though our study identifies OFC value encoding neurons in the human OFC (that is also present in the amygdala and hippocampus) as well, the distinct contributions of these ensembles to decision making and how they evolve throughout time requires further investigation.

## Methods

### Study Participants

9 participants with drug-resistant epilepsy (4 female, 5 male) undergoing depth electrode placement for localization of seizure foci at UCLA were included in the study. All participants provided written, informed consent to participate in research under the approval of the UCLA Medical Institutional Review Board. Participants were told they were free to withdraw consent and discontinue participation at any time. Clinical consideration for surgery was made by a multidisciplinary team of neurosurgeons, neurologists, and neuropsychologists and was independent of the research study. Pre-determined clinical criteria guided placement of Behnke–Fried electrodes (Adtech Medical, Racine WI) in each participant. Electrodes were implanted stereotactically with the aid of digital subtraction computed tomography (CT) angiography and magnetic resonance imaging (MRI). Each Behnke–Fried macro–micro depth electrode contained at least seven macroelectrode contacts (1.5 mm in diameter) spaced 1.5–3.5 mm apart along the shaft, and a Behnke–Fried inner platinum-iridium microwire bundle at the distal end of the electrode (California Fine Wire, Grover Beach, CA; Fried, et al 1999).

### Delay Discounting Task

Participants were presented with choices between $10 available after a specified delay (0, 2, 60, 180, or 365 days) and a smaller amount (< $10) available immediately. For example, the participant could be presented with the choice: “would you rather have $10 in 30 days or $2 now?” The task ran on a laptop using Matlab with Psychtoolbox extensions. Trials started with a fixation cross for 1000 ms after which participants indicated their preference by pressing a button on a Cedrus Response Pad (Response Pad RB-844, Cedrus Corporation, San Pedro, CA 90734, USA). After the participant selected their preferred option, there was a 1000ms inter-trial interval before the fixation cross for the next trial was displayed on the screen. The position of the options on the screen (i.e., the immediate and delayed reward) was counterbalanced across trials. The task utilized an adjusting amount procedure (adjusting the immediate amount in increments of ± $0.50) to derive indifference points between the delayed-standard and immediate-adjusting options for each of the five delays assessed. An indifference point reflected the smallest amount of money an individual chose to receive immediately instead of the delayed standard amount ($10) at the specific delay. Trials continued until indifference points were found for all delays and a discounting quotient, *k*, was calculated. The number of selections that a participant made for immediate vs. delayed rewards and the participant-specific discounting *k* are in Supplementary Table 2.

### Calculation of Participant-Specific Discounting Rate

The reduction in subjective value (SV) of temporally delayed rewards was modeled by the hyperbolic function

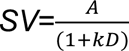

where *A* is the objective amount of the reward (fixed at $10 in our experiment), *D* is the variable delay at which *A* will be administered (2, 60, 180, or 365 days) and *k* is the subject-specific discount rate (Peters 2011). To calculate *k*, the hyperbolic model was fit to indifference points using non-linear least squares via lsqcurvefit.m in Matlab. Smaller values of *k* indicate greater tolerance for extended delays, while larger values of *k* indicate impatience and a refusal to wait for larger rewards to be disbursed in the future.

### Electrode localization

The anatomical location of each electrode was determined by co-registering a high-resolution post-operative computed tomography (CT) scan to a pre-operative whole brain magnetic resonance imaging (T1-weighted sequences). The electrodes were localized by thresholding the raw CT image and calculating the unweighted mass center of each electrode and microelectrode bundle. The preimplantation three-dimensional T1 MR scan was processed using FreeSurfer to segment the white matter, deep gray matter structures and cortex. It was also processed to parcellate the neocortex according to gyral anatomy. Macro- and micro-electrode contacts were then attributed to a cortical region according to automated parcellation in FreeSurfer. Recording contacts were in a variety of regions. Recording channels inside the amygdala, hippocampus and orbitofrontal cortex were included for further analyses. We warped the aligned electrodes onto a standard brain template (using a Montreal Neurological Institute (MNI) template) to facilitate group-level visualization. The MNI reconstruction was performed for visualization purposes only, and electrode localizations were always determined in each patient’s native MRI space.

### Electrophysiology data acquisition

Each depth electrode terminated in eight 40-mm platinum-iridium microwires from which extracellular signals were continuously recorded (referenced locally to a ninth low-impedance microwire). Data were recorded using either Neuralynx (40 kHz sampling rate) or Blackrock (30 kHz sampling rate) data acquisition systems.

### Offline spike sorting and pre processing

Neuronal clusters were identified using the ‘Wave Clus’ software package. As described previously, extracellular recordings were high pass filtered above 300 Hz and a threshold of 5 s.d. above the median noise level was computed (Quiroga et al 2004; Suthana 2015). Detected events were clustered (or categorized as noise) using automatic superparamagnetic clustering of wavelet coefficients, followed by manual refinement based on the consistency of spike waveforms and inter-spike interval distributions. 193 neural clusters (9 patients) were identified by ‘Wave Clus’, utilizing amplitude thresholding and the wavelet transform to implement superparamagnetic clustering. Furthermore, multiunits and single units were classified based on (i) spike shape and variance; (ii) presence of a refractory period (less than 1% spikes with less than 3 ms ISI; (iii) the ratio between the spike peak value and the noise level; and (iv) the ISI distribution of each cluster. There were few multi units (n=9), and restricting the results to single units did not make a significant difference. The presented results therefore incorporate data from both multi units and single units (Rutishauser 2006). To allow for consistency across trials, spike trains were aligned to decision onset and downsampled to 1 kHz. A continuous spike-density function was then calculated using a 100ms standard deviation Gaussian smoothing kernel to estimate firing rate. We used these firing rates to assess task-evoked events.

### Relationship Between Decision Difficulty and Reaction time

Trial difficulty was approximated via the difference in the value of the immediate offer (some amount less than $10) and the subjective value of the delayed offer. Subjective value was calculated using the participant’s individual discounting quotient (*k*), the offer amount ($10), and the delay (2, 60, 180, or 365 days) at which the offer would be withheld (see Calculation of Participant-Specific Discounting Rate). The trial-by-trial difficulty and the corresponding reaction times across participants were fit to a gaussian function of the form,

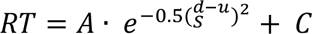

where RT is the reaction time, *A* is the peak height of the function above baseline, *d* is the trial difficulty (i.e. difference between the immediate offer amount and the subjective value of the delayed offer), *μ* is the location of the peak, *S* reflects the width of the function and *C* a constant (Hoffman 2008). In this way, we determined whether a function previously used to model the relationship between reaction time and decision difficulty would also be fit by our data. Data was fit to the function using fit.m in Matlab.

### Identification of Task Variable Encoding Units

For each unit, the decision aligned spike density function for each trial was further downsampled by taking the average activity across non-overlapping 100ms bins from -2 to 0.5 seconds relative to decision onset. Predictor variables (features) for a model were then built by concatenating 5 adjacent bins (average spike density in 100ms) beginning at -2 seconds relative to decision onset. A model was then trained to predict a trial variable (for decision classifiers: immediate vs. delayed choice; for difficulty classifiers: easy vs. hard trial) using 70% of a subject’s trials and then tested on the remaining unseen 30%. Unique training and test sets at identical timepoints were generated across 200 cross validations. Chance distributions were generated by shuffling trial labels. To characterize model accuracy across time, the 5 adjacent bins were shifted every 100ms until 0.5 seconds after decision onset. Models were implemented using LogisticRegressionCV of the sci-kit learn toolbox in python with L2 regularization (Pedegrosa, 2011).

Permutation testing was used to assess whether the experimental decoder performed significantly better than the chance decoder at each 500 ms window by shuffling chance and experimental decoder performances 1000 times to generate a p value. A unit was included in subsequent single unit and population decoding analyses if decoding was significant under a Bonferroni adjusted (with respect to the number of temporally distinct permutation tests performed prior to decision onset) p-value (*p=*0.05/20 for 20 total pre-decision windows). The average accuracy of all variable-decoding units in the OFC, amygdala and hippocampus are included in Figure 2.

### Comparison of Single Unit Decoding Dynamics

The maximum decoding accuracy, and the time at which it occurred, was variable across units within a region and varied based on the variable being decoded (Figure 3; Supplementary Figure 2). A linear mixed effects model was used to determine if there were significant differences in average peak decoding performance and timing within a region based on the variable being decoded. Within the LMM, participant and unit identity were treated as random effects. The restricted maximum likelihood method was used to estimate an effect of task variable (difficulty decoding: easy vs. hard; choice decoding: immediate vs. delayed). In this way, the model determined whether the estimated difference in average peak accuracy or average peak accuracy time for choice and difficulty decoding within a region was significantly different from zero.

### Population Decoding Analyses

The collective decoding capacity of units was assessed by decoding from the population of recorded units. Here, model features consisted of the average firing rate of each unit in a 500 ms window. All units (separately for each region) that exhibited significant choice/difficulty decoding were used as model features. The model was trained on 50% of the data and was tested on the remaining 50%.

Training and test set construction began by splitting each cell’s activity (at a specific point in time) during each condition (i.e., immediate vs. delayed when decoding choice) into half (one half used for training, and the other for testing). Because participants had a unique preference for immediate vs. delayed rewards, they had a different number of exposures to each condition with respect to choice (immediate vs. delayed) and difficulty (easy vs. hard) (Supplementary Table 2). Thus, the training and test set size was bounded by the minimum # of trials seen for each condition across subjects. For example, the minimum of all delayed and immediate choice trials across participants was 10 and, thus, 5 immediate and delayed trials (10 total) were used for testing. To maximize the amount of data that the model was exposed to prior to testing, model features for each trial in the training data was constructed by randomly selecting a cell’s activity (feature) during a trial (that was not used in testing) during a specific condition. Specifically, a model saw features N_1_, N_2_, N_3_, …N_n_ where n = number of regional (i.e. OFC, HPC, amygdala), choice, or difficulty selective cells. For a training example for a condition of interest (i.e. immediate or delayed when decoding choice), N_1_ consisted of cell 1’s firing rate randomly selected from a pool containing that cell’s activity during unique instances of that condition. Features N_1_ to N_n_ consisted of unique cells during identical conditions across training examples. As in single unit decoding analyses, unique training and test sets at identical timepoints were generated across 200 cross validations. Model efficacy was assessed by comparing performance of the experimental decoder to chance decoders that were generated by training a decoder on the same training data, but with shuffled trial labels. All population models were implemented using LogisticRegressionCV of the sci-kit learn toolbox in python with L2 regularization (Pedegrosa, 2011).

We used two approaches to assessing population encoding of a variable of interest. In the first approach, a fixed, an identical time window was used across all units. Given that units varied based on the timing of when they were most predictive of decision/decision difficulty, we also built models using activity centered at the point at which an individual unit was most predictive of the variable of interest. Across all regions, utilization of activity centered at each unit’s most predictive temporal window led to a higher mean accuracy than the greatest mean accuracy observed in the fixed temporal window approach (Supplementary Figure 6).

### Fast vs. Slow Discounting Models

We divided participants into two groups based on their discounting behaviors. Slow discounters were willing to accept temporal delays in exchange for greater monetary rewards and had an average *k* value of 0.0015 +/- 0.0005 (s.e.m.) and an average reaction time of 3.99 +/- 0.10 (s.e.m.) seconds. Fast discounters typically chose the immediately available option and had an average *k* value of 0.1739 +/- 0.084 and an average reaction time of 5.42 +/- 0.22 (s.e.m.) seconds. We used two approaches to approximate differences in trial variable representations between participants that preferred immediate rewards (fast discounters) vs. those that were relatively agnostic to temporal delays (slow discounters). In both approaches, we used the peak-population decoding strategy discussed in *Population Decoding Analyses*. In the first approach, we compared regional population decoders built from units from only fast discounters (“Fast Only”) to regional decoders built from an equal number of units randomly sampled from all participants (“All (fixed)”). Both “Fast Only” and “All (fixed)” contained N_1_ to N_Fs_ features where Fs = number of unique units isolated across participants with a tendency to rapidly discount temporally delayed rewards (Supplementary Table 2). The “All (fixed)” model represented an approximation of the discounting behavior agnostic model that included all participants described in *Population Decoding Analyses*, but with a number of features identical to that used in the “Fast Only” model. A total of Fs units/features were randomly selected across all participants to construct the All (fixed) model since decoder performance may increase with more units. The expanded training set approach described in *Population Decoding Analyses* was also used in both models here. In the second approach, we compared regional population decoders built from units from only slow discounters to regional population decoders built from an equal number of units from fast discounters.

### Fast vs. Slow Single Unit Characteristics

To determine if participant behavior is related to the level of variable encoding at the single unit level, we compared the average single unit encoding of choice/difficulty (see Identification of Task Variable Encoding Units; Supplementary Figure 8) in fast vs. slow discounters using a linear mixed effects model. As in previous single unit analyses (see Comparison of Single Unit Decoding Dynamics), participant and unit identity were treated as random effects. The restricted maximum likelihood method was used to estimate an effect of trial variable. To determine if the time at which single unit activity was most predictive of choice/difficulty was different in fast vs. slow discounters we used the same linear mixed effects model approach using peak decoding accuracy time as the predictor variable (Supplementary Figure 9).

---

**Supplementary Figure 1.**
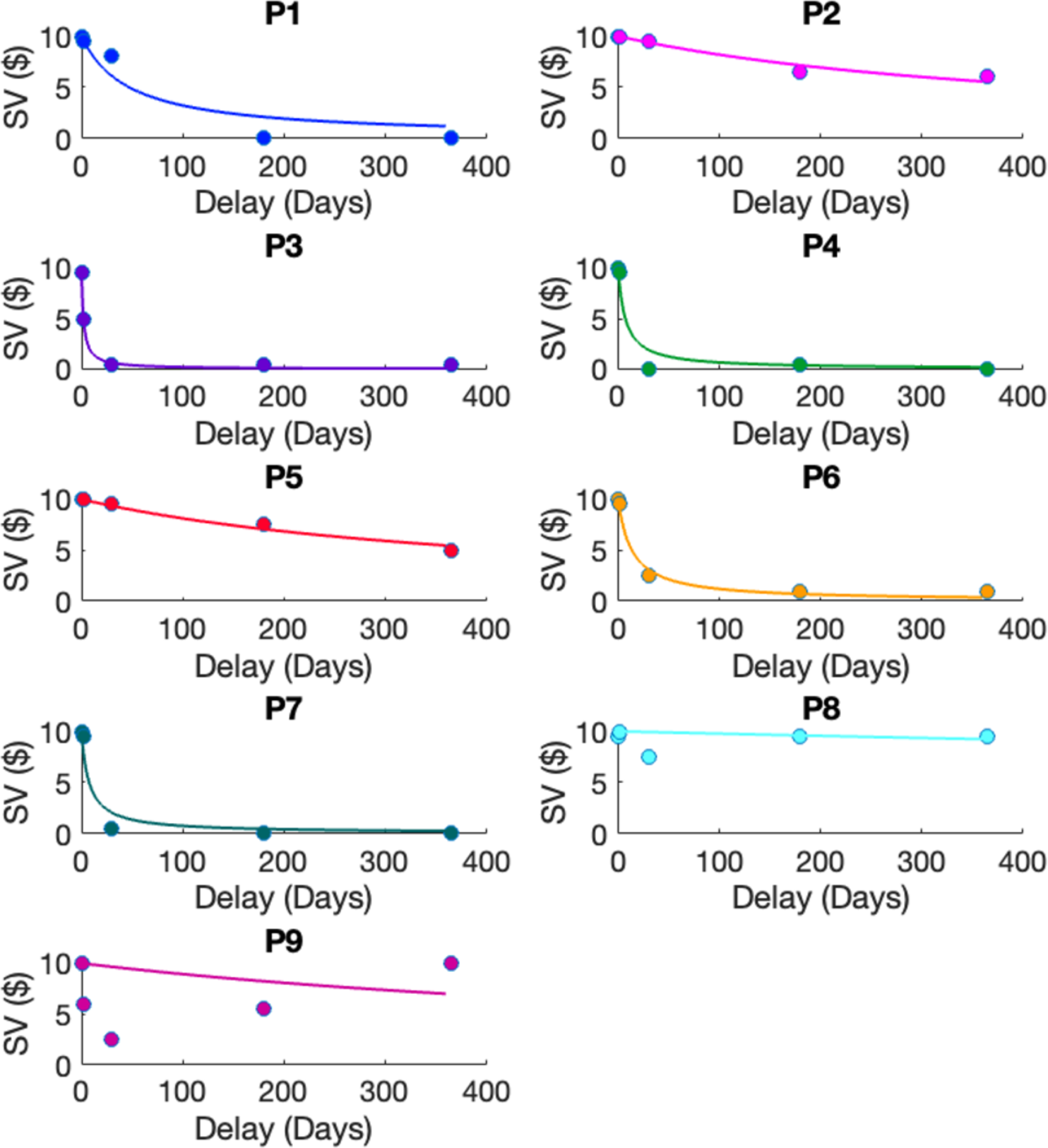
Discounting curves for each participant (solid line) fit to indifference points (point at which at which y axis $ value was similarly valued to $10 at x axis delay; dots). Discounting curves were fit to indifference points using a one-parameter hyperbolic model (see Calculation of Participant-Specific Discounting Rate, Methods). Discounting curves with steeper slopes indicate that an individual rapidly reduces the subjective value (y axis) of a large, potential reward with increasing delay (x axis). Colors correspond to those used in Figure 1.

**Supplementary Figure 2.**
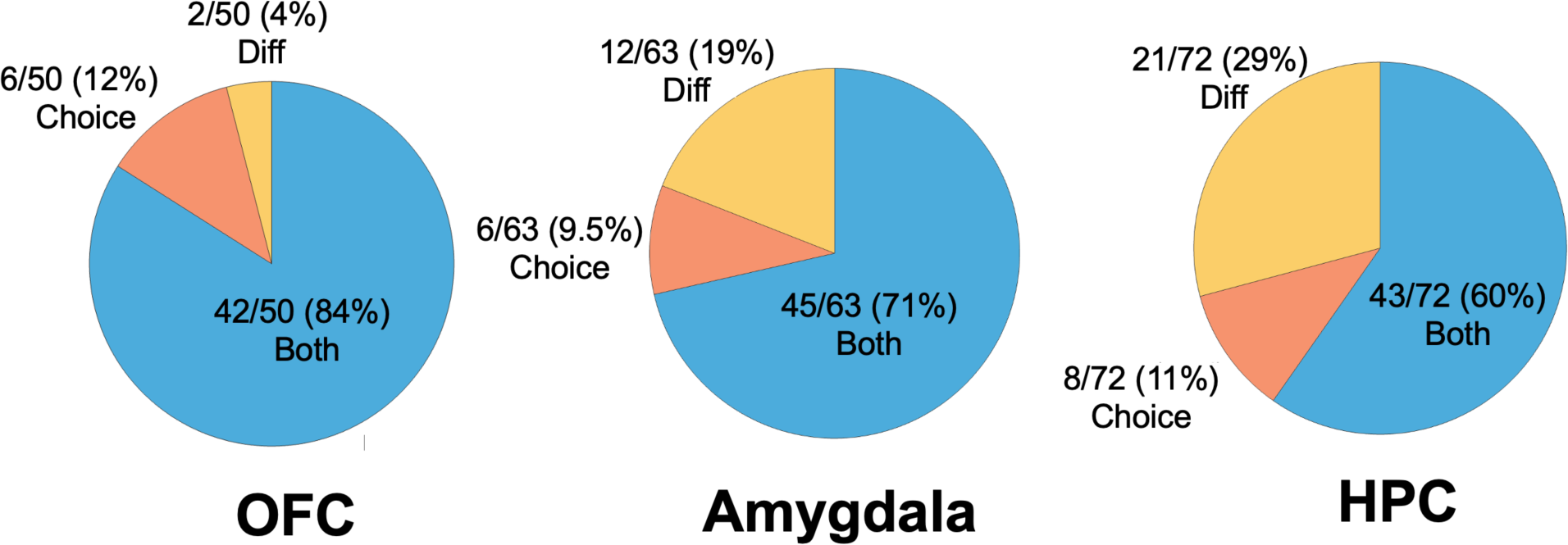
Unit Selectivity. Number of units that significantly predicted choice, difficulty, or both over the number of units that predicted choice or difficulty within the orbitofrontal cortex (OFC), amygdala or hippocampus (HPC).

**Supplementary Figure 3.**
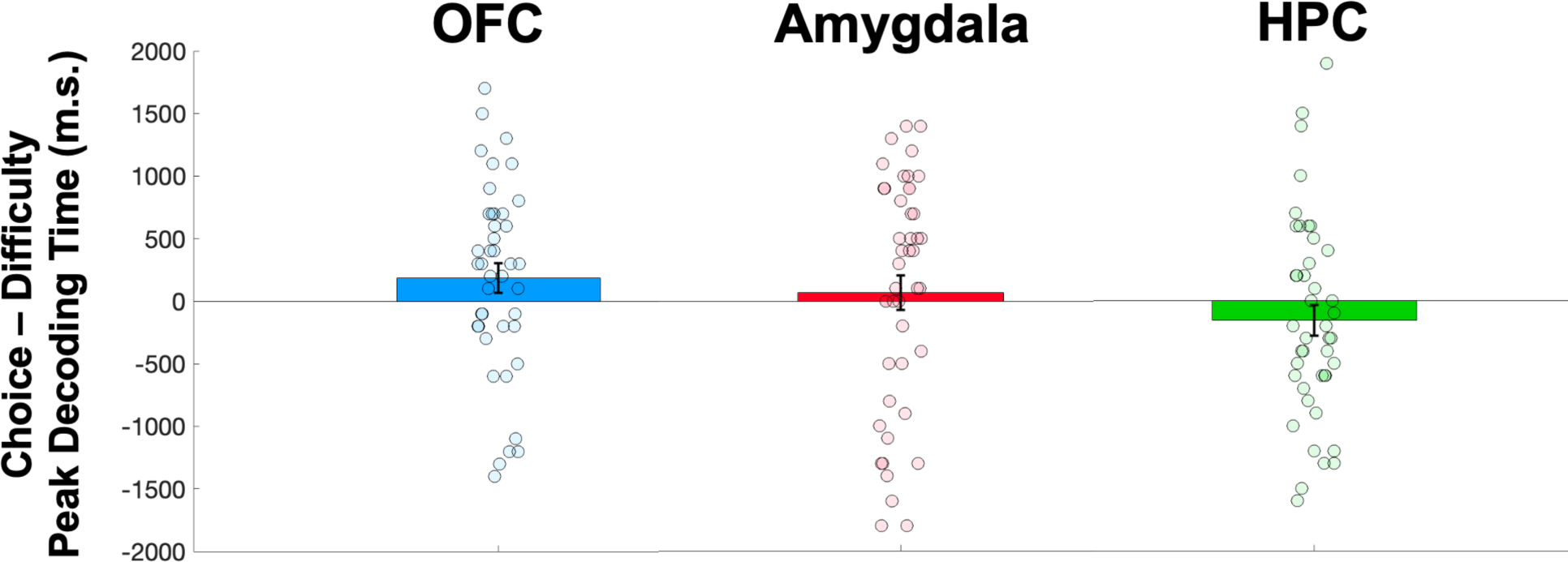
Peak Decoding Time Differences in Units that Encoded Choice and Difficulty. Average millisecond (m.s.) difference in peak (time at which decoding accuracy prior to decision onset was highest prior to choice; Methods) decoding of choice and difficulty for individual units across n = 9 participants that decoded both choice and difficulty (Supplementary Figure 1) in the orbitofrontal cortex (OFC; blue; n = 42 units, choice mean = -952.38 ms +/- 14.90 s.e.m; difficulty mean = - 1138.10 ms +/- 12.92 s.e.m; p = .073), amygdala (red; n = 45 units, choice mean = - 1000.00 ms +/- 13.25 s.e.m; difficulty mean = -1068.90 ms +/- 13.12 s.e.m; p = .604) and the hippocampus (HPC; green; n = 43 units, choice mean = -1116.30 ms +/- 13.80 s.e.m; difficulty mean = -960.47 ms +/- 14.68 s.e.m; p = .159). Positive values indicate peak decoding accuracy of difficulty occurring before peak decoding time of choice and vice versa for negative values. Wilcoxon signed rank test of choice vs. difficulty peak decoding times did not achieve significance in any region (Methods).

**Supplementary Figure 4.**
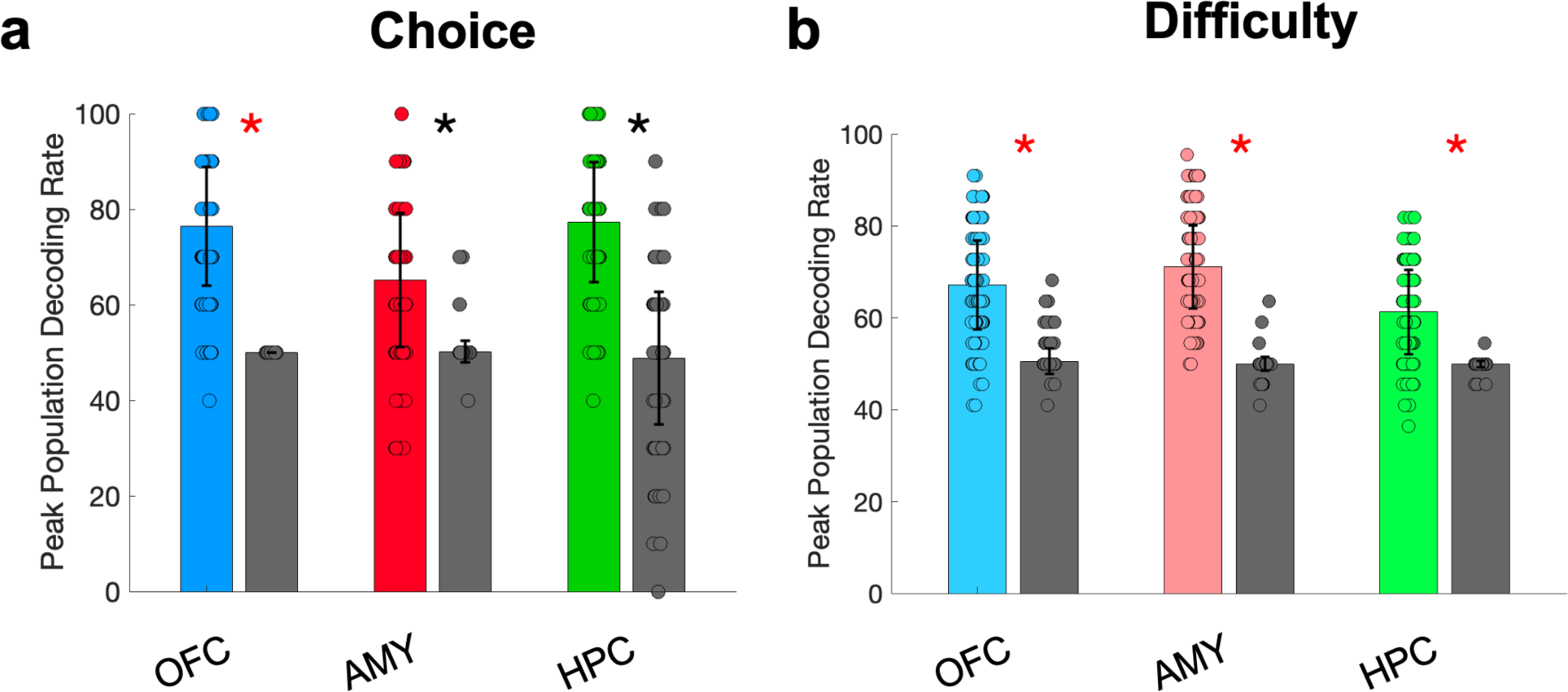
Multiregional Choice and Difficulty Peak Population Decoding. **a)** Average accuracy +/- standard deviation (s.d) in the orbitofrontal cortex (OFC; blue), amygdala [red] and hippocampus [HPC; green] of a peak population decoder trained to predict trial choice prior to decision onset (Methods). Gray bar represents the corresponding average accuracy of a chance decoder trained on identical data, but with shuffled labels (Methods). Asterisks indicate above chance decoding. black * = p <.05, red * = p <.01. **b)** same as **a** but for a decoder trained to predict decision difficulty prior to choice (Methods).

**Supplementary Figure 5.**
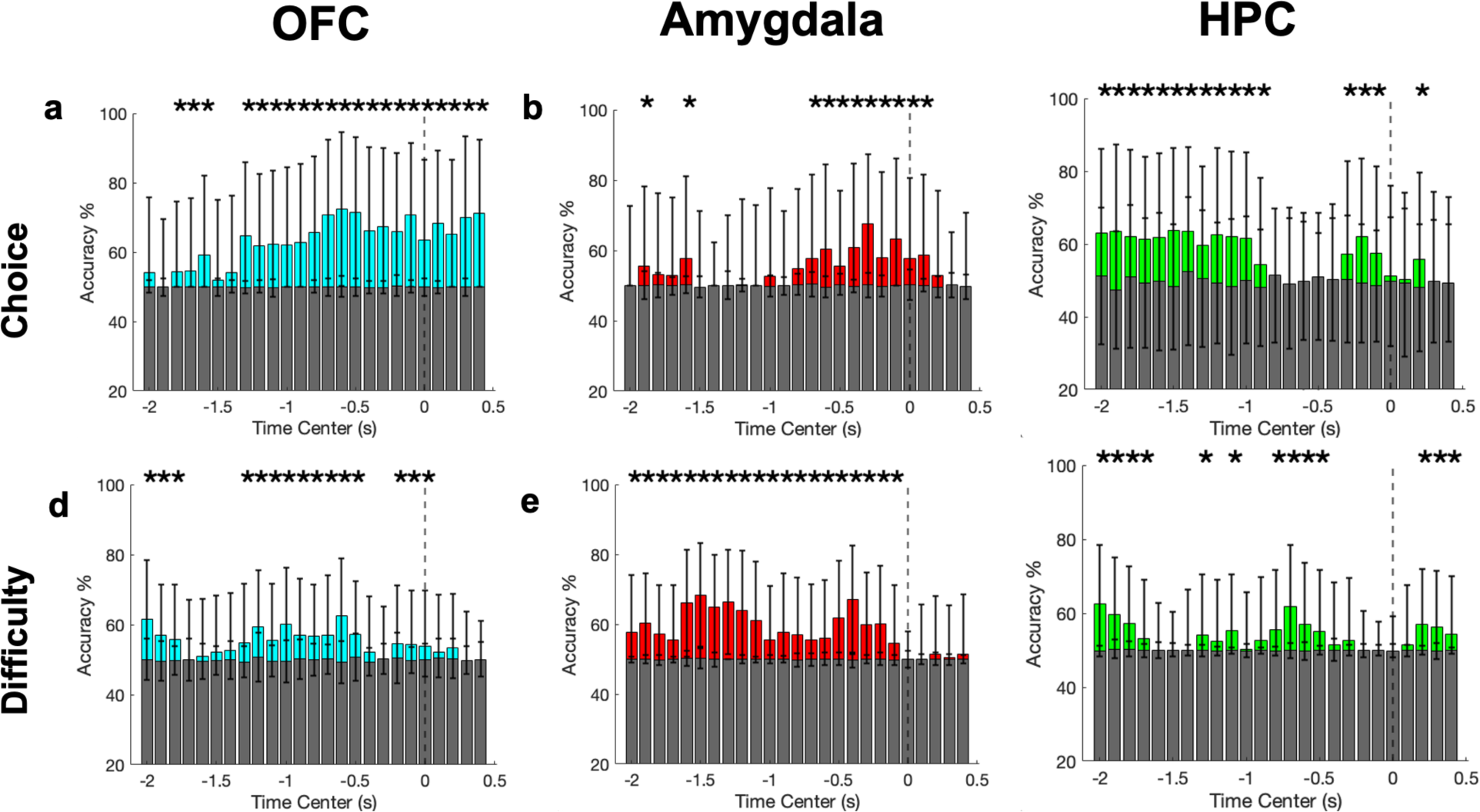
Multiregional Choice and Difficulty Peak Population Decoding. **a)** Average accuracy +/- standard deviation [s.d] in the orbitofrontal cortex [OFC; blue], amygdala [red] and hippocampus [HPC; green] of a population decoder trained to predict trial choice (top row; **a-c)** or difficulty (bottom row; **d-f)** prior to decision onset (Methods). Gray bar represents the corresponding average accuracy of a chance decoder trained on identical data, but with shuffled labels (Methods). Asterisks indicate above chance decoding. * = p <.05 using permutation testing.

**Supplementary Figure 6.**
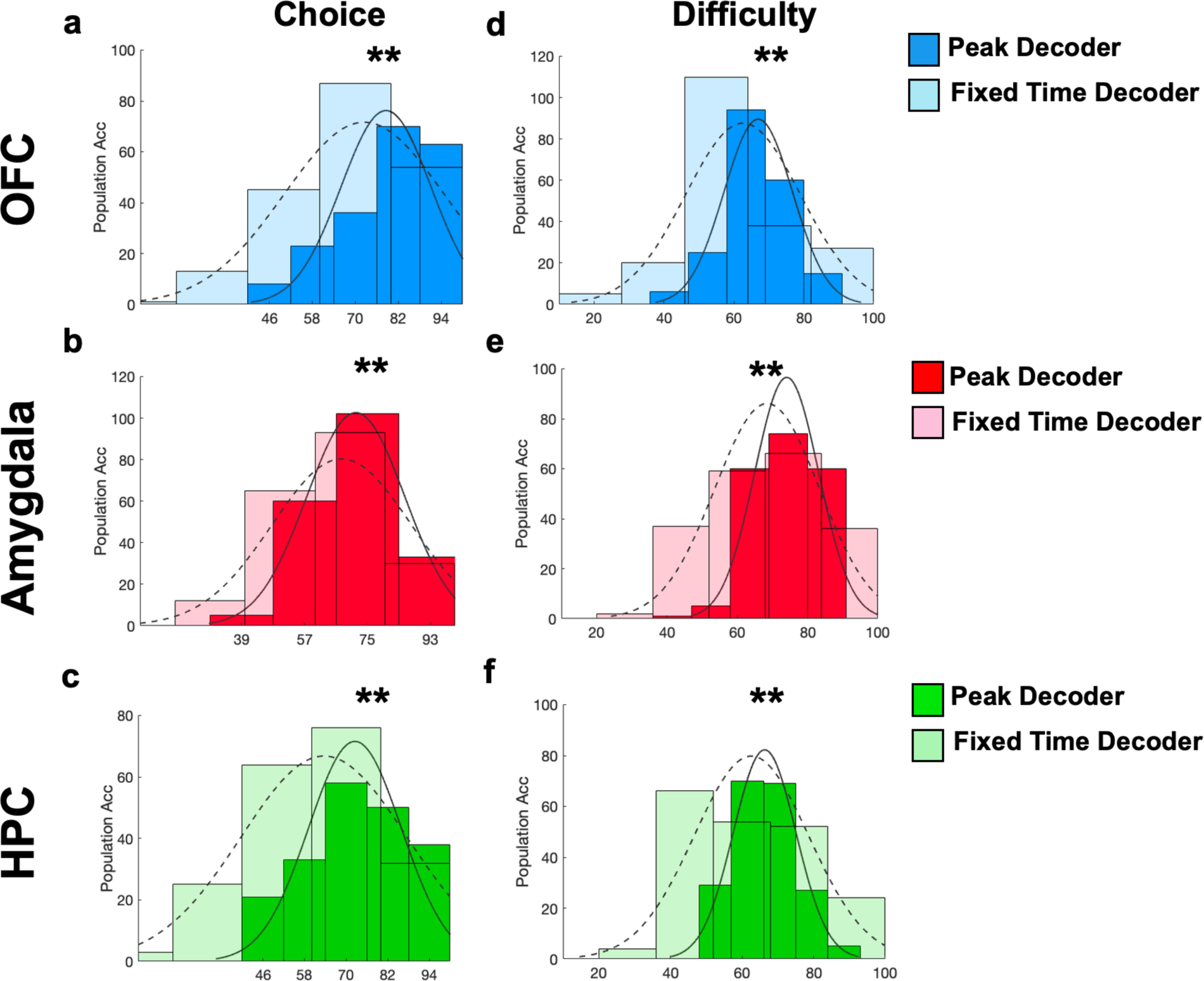
Improved Population Decoding with Peak Accuracy Decoding Approach. **a-c)** Histogram of decoding performance across 200 cross validations from a model built using average activity from units (features) in 500 ms centered where decoding accuracy peaked (darker shade of all colors; Peak Decoder; Methods) vs. a model using activity centered at the same time point across units when collective accuracy was highest (lighter shade of all colors; Fixed Time Decoder; Methods) during decoding of choice (left) and difficulty (right). Units were isolated from **a, d)** the orbitofrontal cortex [OFC], **b, e)** the amygdala and **c, f)** the hippocampus (HPC). * = p<.05, ** = p<.01 using Wilcoxon rank sum(Methods). **a)** OFC choice mean +/- standard deviation (s.d.) = Peak: 78.6 +/- 12.56; Fixed: 72.50 +/- 22.27. **d)** OFC difficulty mean +/- s.d = Peak: 66.97 +/- 9.79; Fixed: 62.63 +/- 16.38. **b)** Amygdala choice mean +/- s.d = Peak: 71.7 +/- 13.99; Fixed: 67.63 +/- 19.86. **e)** Amygdala difficulty mean +/- s.d = Peak: 74.06 +/- 9; Fixed: 68.50 +/-14.83. **c)** HPC choice mean +/- s.d = Peak: 72.55+/- 13.38; **f)** Fixed: 63.250 +/- 24.02. HPC difficulty mean +/- s.d = Peak: 66.32 +/- 8.73;Fixed: 62.65 +/- 15.99.

**Supplementary Figure 7.**
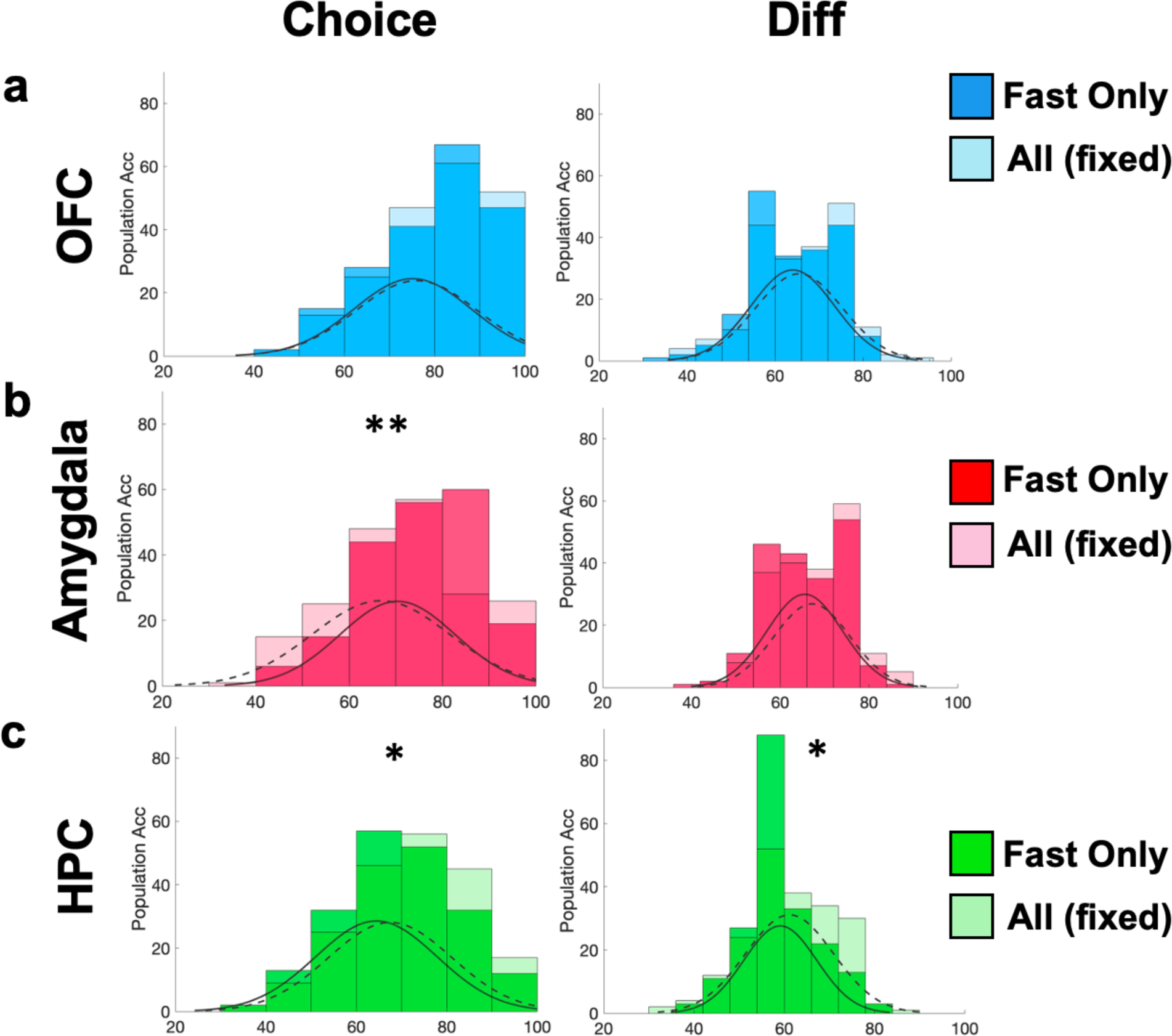
Altered Decoding in Fast Discounter Model. **a-c)** Histogram of decoding performance across 200 cross validations from a model built using average activity from units (features) from participants with fast discounting rates (darker shade of all colors; fast only) vs. a model built using a matched number of randomly selected units from all participants (lighter shade of all colors; All (fixed)) during decoding of choice (left) and difficulty (diff; right). Units were located in: **a)** the orbitofrontal cortex (OFC, fast only choice mean +/- standard deviation (s.d.) = 75.15% +/- 13.03, all (fixed) choice mean +/- s.d. = 76.05% +/- 13.37, p = .451; fast only diff mean +/- s.d. = 64.03 % +/- 9.48, all (fixed) diff mean +/- s.d. = 65.45% +/- 9.90, p = .130), **b)** the amygdala (fast only choice mean +/- s.d. = 70.40% +/- 12.35, all (fixed) choice mean +/- s.d. = 66.85% +/- 14.72, p = .009; fast only diff mean +/- s.d. = 65.55% +/- 8.52, all (fixed) diff mean +/- s.d. = 67.18% +/- 8.61, p = .054) and **c)** the hippocampus (HPC, fast only choice mean +/- s.d. = 59.13% +/- 8.08; all (fixed) choice mean +/- s.d. = 61.18 % +/- 9.74; p = .024). * = p<.05, ** = p<.01 using permutation testing (Methods).

**Supplementary Figure 8.**
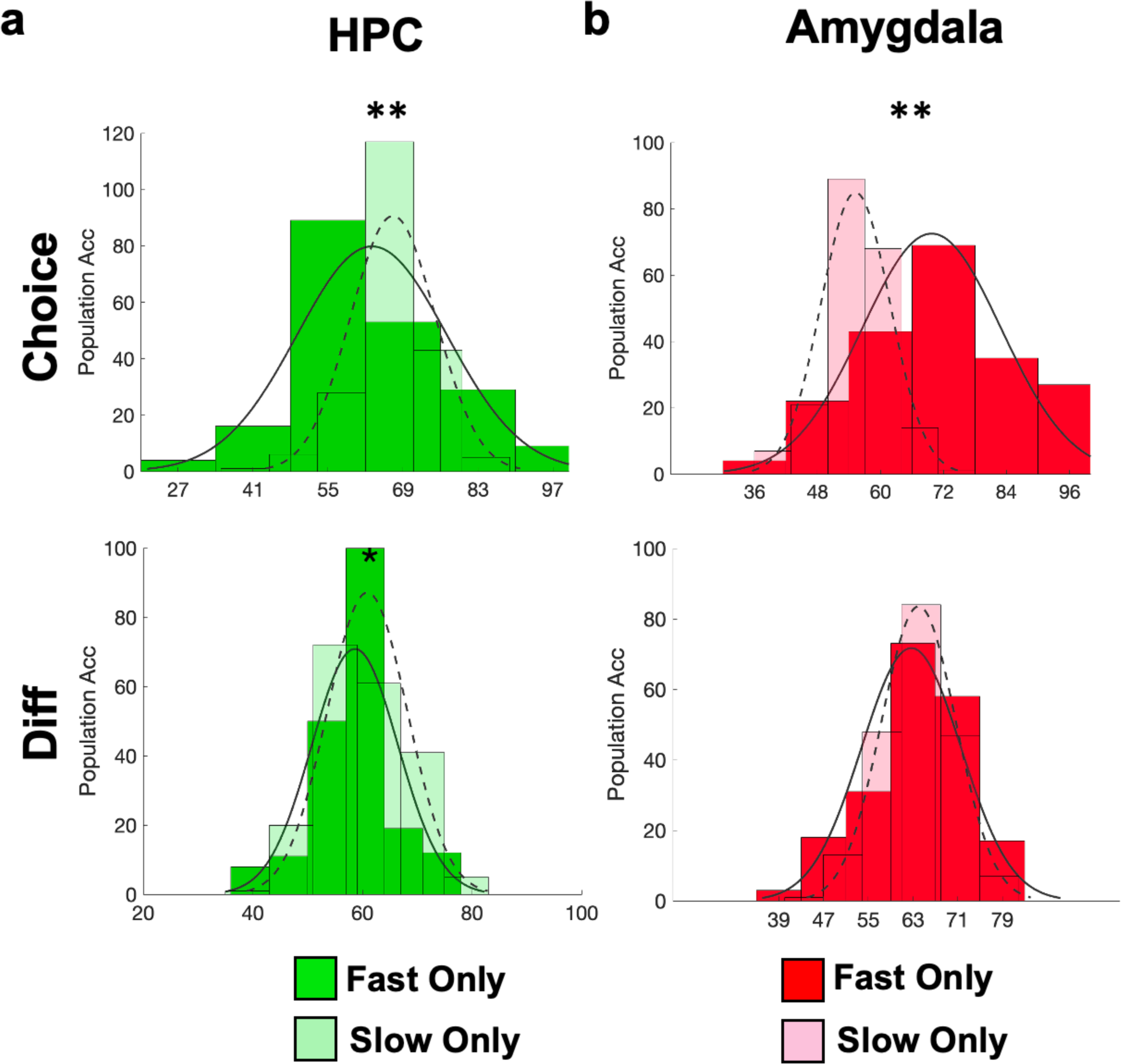
Differential Performance in Slow and Fast Discounter Models. **a-b)** Histogram of decoding performance across 200 cross validations from a model built using average activity from units (features) from participants with fast discounting rates (darker shade of red/green; Fast Only) vs. a model built using a matched number of units from all participants with slow discounting rates (lighter shade of red/green; Slow Only) during decoding of choice (top row) and difficulty (right). Units were isolated from **a)** the hippocampus (HPC, fast only choice mean +/- standard deviation (s.d.) = 63.20% +/- 13.99, slow only choice mean +/- s.d. = 67.13% +/- 7.92, p < .001; fast only diff mean +/- s.d. = 58.6% +/- 7.88, slow only diff mean +/- s.d. = 60.90 % +/- 7.32, p = .030) and **b)** the amygdala (fast only choice mean +/- s.d. = 69.75 % +/- 13.20, slow only choice mean +/- s.d. = 55.25% +/- 6.57, p < .001; fast only diff mean +/- s.d. = 62.70% +/- 8.89, slow only diff mean +/- s.d. = 64.03% +/- 6.68, p =.247). * = p<.05, ** = p<.01 using Wilcoxon rank sum (Methods)

**Supplementary Figure 9.**
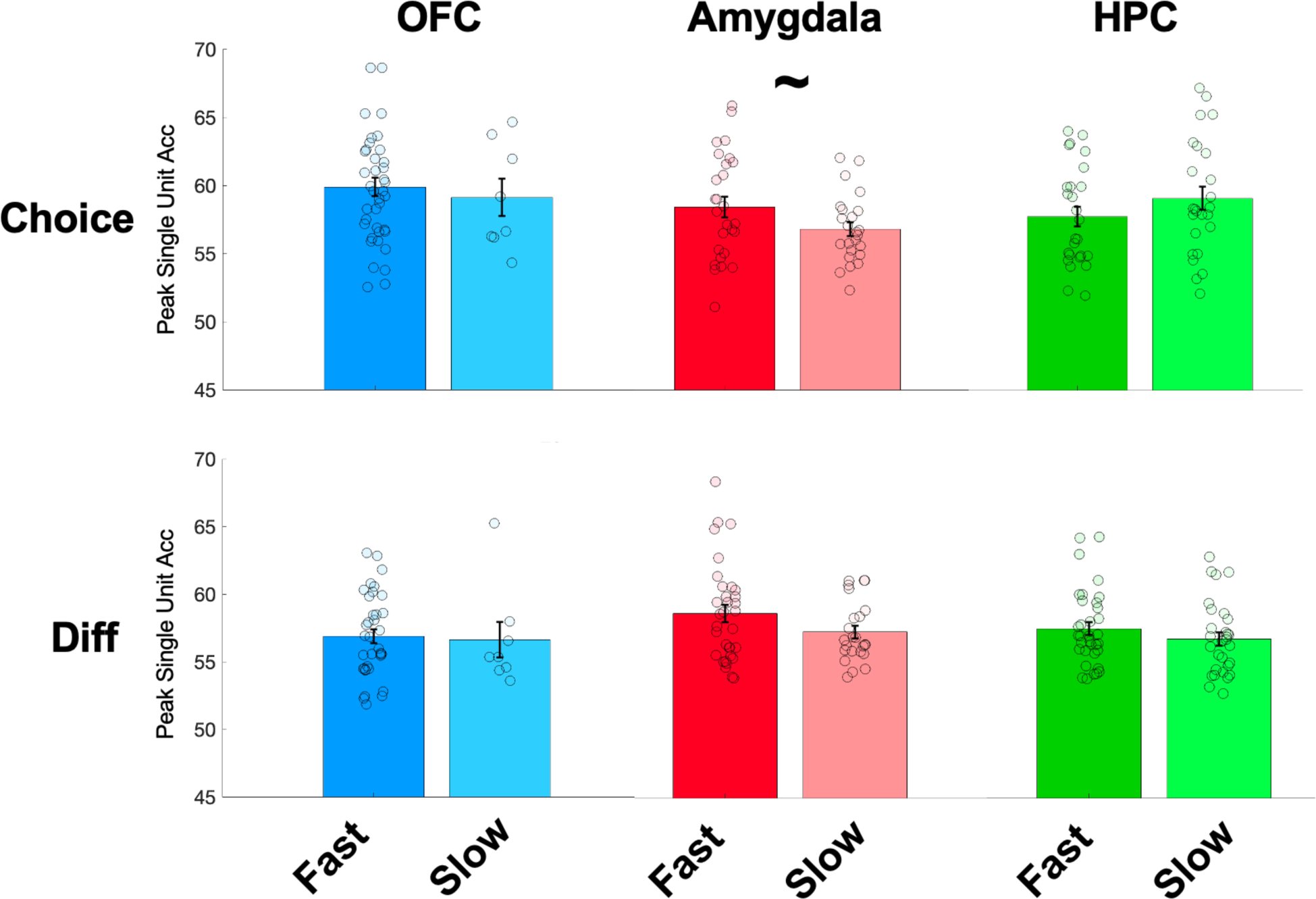
Single Unit Peak Decoding Accuracy In Slow and Fast Discounters. Average peak single unit choice (top row) and difficulty (Diff; bottom row) decoding accuracy in slow and fast discounters (Methods) in the orbitofrontal cortex (OFC; blue) amygdala (red) and hippocampus (HPC; green). ∼ = p <.1 using a linear mixed effects model (Methods). OFC average fast discounter unit peak choice decoding accuracy +/- standard error of the mean (s.e.m.) = 59.88% +/- .68, average slow discounter unit peak choice decoding accuracy +/- s.e.m. = 59.12% +/- 1.37; p = .359. OFC average fast discounter unit peak diff decoding accuracy +/- s.e.m.= 56.88% +/- .51, average slow discounter unit peak diff decoding accuracy +/- s.e.m.= 56.62% +/- 1.32 ; p = .885. Amygdala average fast discounter unit peak choice decoding accuracy +/- s.e.m. = 58.56% +/- .75, average slow discounter unit peak choice decoding accuracy +/- s.e.m. = 56.92% +/- .51; p = .085. Amygdala average fast discounter unit peak diff decoding accuracy +/- s.e.m.= 58.23% +/- .61, average slow discounter unit peak diff decoding accuracy +/- s.e.m.= 56.90% +/- .44; p = .215. HPC average fast discounter unit peak choice decoding accuracy +/- s.e.m. = 57.90% +/- .73, average slow discounter unit peak choice decoding accuracy +/- s.e.m. = 59.26 +/- .86 ; p = .897. HPC average fast discounter unit peak diff decoding accuracy +/- s.e.m. = 57.12% +/- .46, average slow discounter unit peak diff decoding accuracy +/- s.e.m. = 56.39%+/- .47; p = .367.

**Supplementary Figure 10.**
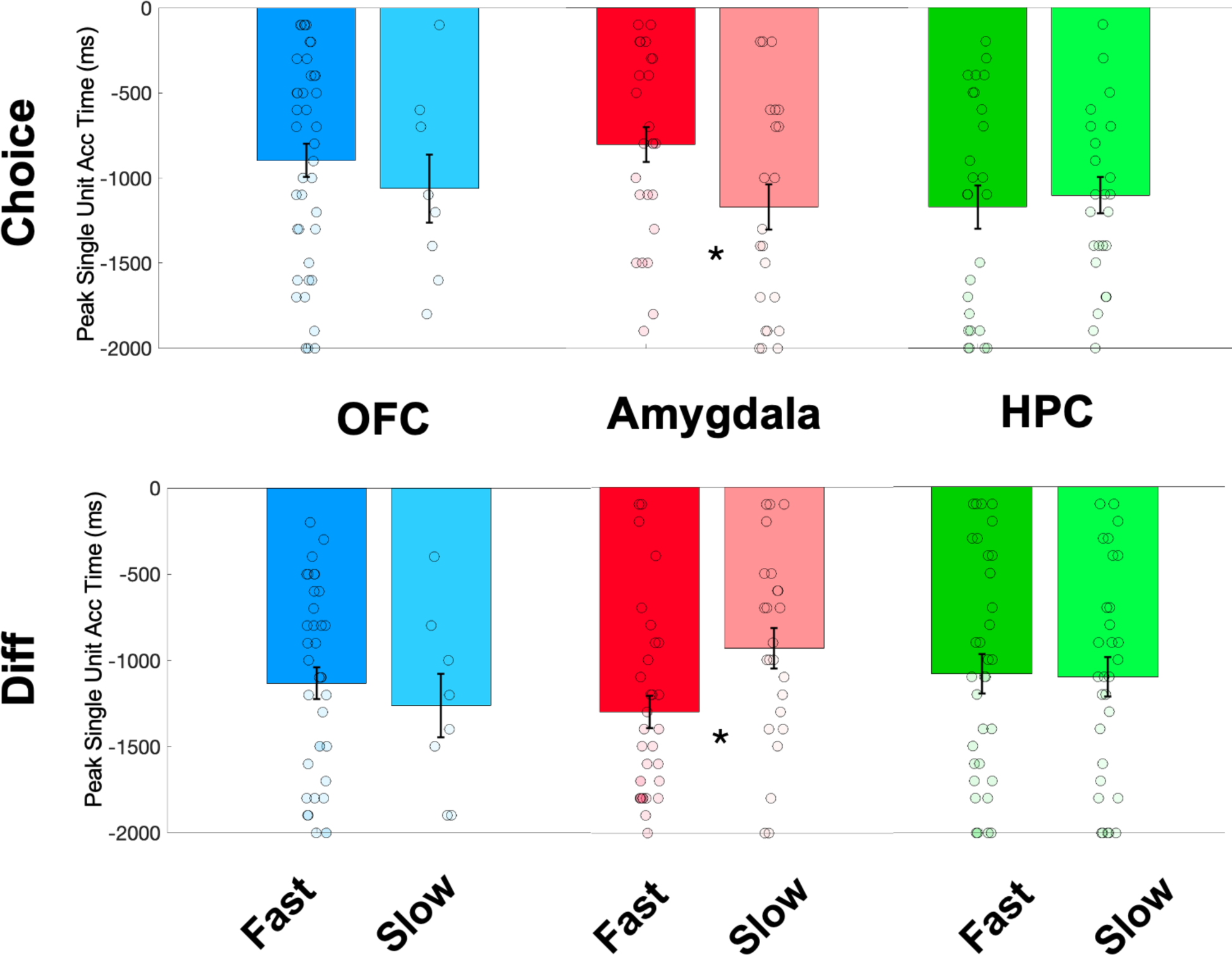
Peak Decoding Time Differences in Slow and Fast Discounters. Average millisecond (m.s.) peak decoding time (time at which decoding accuracy was highest prior to choice; Methods) of choice (top row) and difficulty (Diff; bottom row) for individual units across slow and fast discounters (Methods) in the orbitofrontal cortex (OFC; blue), amygdala (red) and the hippocampus (HPC; green). * = p <.05 using a linear mixed effects model (Methods). OFC average fast discounter unit peak choice decoding time +/- standard error of the mean (s.e.m.) = -897.5 m.s. +/- 199.94, average slow discounter unit peak choice decoding time +/- s.e.m.= -1062.5 m.s +/- 97.96; p =.501. OFC average fast discounter unit peak diff decoding time +/- s.e.m. = -1133.3 m.s. +/- 92.50, average slow discounter unit peak diff decoding time +/- s.e.m. = -1262.5 m.s. +/- 185.10 ; p =. 551. Amygdala average fast discounter unit peak choice decoding time +/- s.e.m. = -803.70 m.s. +/- 102.04, average slow discounter unit peak choice decoding time +/- s.e.m. = -1170.8 m.s. +/- 132.89 ; p = .031. Amygdala average fast discounter unit peak diff decoding time +/- s.e.m. = - 1300.00 m.s. +/- 94.15, average slow discounter unit peak diff decoding time +/- s.e.m. = -933.33 m.s. +/- 115.88; p = .016. HPC average fast discounter unit peak choice decoding time +/- s.e.m.= -1173.10 m.s +/- 126.44, average slow discounter unit peak choice decoding time +/- s.e.m. = -1104.00 m.s +/- 17.31; p = .679. HPC average fast discounter unit peak diff decoding time +/- s.e.m. = -1082.40 m.s. +/- 113.17, average slow discounter unit peak diff decoding time +/- s.e.m. = -1100 m.s. +/- 114.47; p = .959.

**Supplementary Table 1.** Participant Electrode Locations. The MNI coordinates and associated brain regions of microelectrode bundles for all included participants

|  |  | x | y | z | Participant ID |
| --- | --- | --- | --- | --- | --- |
| Amygdala | Left | -21 | -4 | -15 | P1 |
|  |  | -20 | 1 | -26 | P4 |
|  |  | -18 | -5 | -22 | P2 |
|  |  | -18 | -7 | -23 | P5 |
|  |  | -15 | -8 | -20 | P6 |
|  | Right | 16 | -4 | -22 | P6 |
|  |  | 18 | -3 | -21 | P2 |
|  |  | 20 | -2 | -25 | P5 |
|  |  | 21 | -8 | -19 | P1 |
|  |  | 22 | -5 | -27 | P8 |
|  |  | 23 | -1 | -25 | P9 |
|  |  | 29 | -7 | -19 | P7 |
| Hippocampus | Left | -30 | -22 | -14 | P2 |
|  |  | -26 | -18 | -15 | P4 |
|  |  | -25 | -18 | -19 | P3 |
|  |  | -24 | -12 | -20 | P1 |
|  |  | -23 | -19 | -16 | P5 |
|  |  | -23 | -15 | -19 | P7 |
|  |  | -18 | -19 | -16 | P6 |
|  |  | -12 | -40 | -8 | P3 |
|  | Right | 20 | -16 | -25 | P1 |
|  |  | 21 | -14 | -16 | P8 |
|  |  | 21 | -9 | -22 | P9 |
|  |  | 22 | -13 | -22 | P4 |
|  |  | 23 | -23 | -13 | P3 |
|  |  | 25 | -17 | -18 | P5 |
|  |  | 25 | -13 | -23 | P7 |
|  |  | 26 | -22 | -18 | P6 |
| OFC | Left | -7 | 38 | -7 | P2 |
|  |  | -7 | 42 | -4 | P6 |
|  |  | -7 | 52 | -8 | P9 |
|  |  | -6 | 39 | -12 | P8 |
|  |  | -5 | 37 | -1 | P4 |
|  | Right | 1 | 52 | -6 | P9 |
|  |  | 2 | 36 | -9 | P6 |
|  |  | 4 | 45 | -14 | P8 |
|  |  | 9 | 42 | -2 | P5 |

**Supplementary Table 2.** Participant Behaviors. Identification number (ID), age, sex and handedness for each included participant. Corresponding number of trials where a larger delayed option was chosen over a smaller immediate option (“Delayed Choices”) and vice versa (”Immediate Choices”). “Easy” and “Hard” correspond to number of trials where the difference in subjective value between the immediate and delayed offer was >|$1| and <= |$1| respectively. “k” corresponds to each subject’s discounting quotient, which was used to determine whether they were identified as a “Fast” or “Slow” discounter (“Group”; Methods).

| ID | Age | Sex | Handedness | Delayed Choices | Immediate Choices | Easy | Hard | k value | Group |
| --- | --- | --- | --- | --- | --- | --- | --- | --- | --- |
| P1 | 26 | M | R | 31 | 35 | 54 | 25 | 0.0214 | Fast |
| P2 | 47 | M | L | 61 | 49 | 51 | 66 | 0.0022 | Slow |
| P3 | 22 | F | R | 40 | 34 | 42 | 33 | 0.0012 | Slow |
| P4 | 36 | M | R | 10 | 50 | 45 | 19 | 0.5005 | Fast |
| P5 | 29 | F | R | 16 | 52 | 58 | 24 | 0.1414 | Fast |
| P6 | 31 | F | R | 38 | 37 | 33 | 50 | 0.0002 | Slow |
| P7 | 69 | M | R | 53 | 25 | 49 | 45 | 0.0023 | Slow |
| P8 | 25 | F | R | 37 | 40 | 40 | 54 | 0.0739 | Fast |
| P9 | 50 | M | L | 28 | 47 | 63 | 33 | 0.1321 | Fast |

## Notes

### Competing Interest Statement

The authors have declared no competing interest.

